# Camelina (*Camelina sativa* L. Crantz) Crop Performance, Insect Pollinators, and Pollen Dispersal in the Northeastern US

**DOI:** 10.1101/756619

**Authors:** Richard Rizzitello, Chuan-Jie Zhang, Carol Auer

## Abstract

*Camelina sativa* (camelina) is an oilseed crop in the Brassicaceae that has been genetically engineered for the production of biofuels, dietary supplements, and other industrial compounds. Cultivation in North America is both recent and limited, so there are gaps in knowledge regarding yield, weed competition, and pollen-mediated gene flow. For these experiments, camelina ‘SO-40’ was grown for three years without weed control. Spring-sown camelina was harvested at 80-88 days with ∼1200 growing degree days (GDD) with yields of 425-508 kg/hectare. Camelina yields were the same with or without weeds, showing competitive ability in low-management conditions. Crop failure in 2015 was associated with delayed rainfall and above-average temperatures after seeding. Camelina flowers attracted pollinating insects from the Hymenoptera, Diptera, Lepidoptera, and Coleoptera. Hymenoptera included honey bees (*Apis melifera*), mining bees (Andrenidae), sweat bees (Halictidae), bumble bees (*Bombus* spp.) and leaf cutter bees (Megachilidae). Insect visitation on camelina flowers was associated with modest increases in seed yield. Honey bees comprised 28-33% of all pollinators and were shown to carry camelina pollen on their legs. Air sampling showed that wind-blown pollen was present at low concentrations at 9 m beyond the edges of the field. These experiments demonstrated for the first time that camelina pollen dispersal could occur through honey bees or wind, although bee activity would likely be more significant for long-distance gene flow.

## 1. Introduction

Research on *Camelina sativa* (L.) Crantz (camelina) has demonstrated its potential to produce industrial compounds with specific biological, physical, and chemical properties (Bansal and Durrett, 2016; Horn and Benning, 2016; Wurtzel and Kutchan, 2016; Yang et al., 2016). The primary products extracted from camelina seeds have been modified fatty acids for biofuels, bioplastics and dietary supplements (Bansal and Durrett, 2016; Horn and Benning, 2016; Vollmann and Eynck, 2015). Camelina oil has good fuel properties for petroleum-based diesel engines (Bernardo et al., 2003; Yang et al., 2016). It has also been used successfully in military and commercial jets (http:/susoils.com). Algal genes inserted into genetically engineered (GE) camelina have allowed seed production of high concentrations of long-chain polyunsaturated fatty acids eicosapentaenoic acid (EPA) and docosahexaenoic acid (DHA) (Betancor et al., 2015). These fatty acids have economic value as dietary supplements for aquaculture and other applications (Betancor et al., 2015; Petrie et al., 2014; Usher et al., 2015). The development of plant-based omega-3-fatty acids could reduce the harvest of Atlantic menhaden and other fish species that may be in decline (Aaron Kornbluth, Pew Charitable Trusts, personal communication).

Camelina has also generated some excitement as an adaptable, low-input crop. It grows in cool temperate regions with a short lifecycle, less fertilizer and fewer insect pests compared to other Brassica crops (e.g. oilseed rape) (Bansal and Durrett, 2016; Ehrensing and Guy, 2008; Francis and Warwick, 2009; Gugel and Falk, 2006; Jewett, 2013; McVay and Lamb, 2004; Zanetti et al., 2017; Zubr, 1997). While some studies have noted good drought tolerance (Zanetti et al., 2017), other studies have documented crop failure due to drought (Eberle et al., 2015). In addition, camelina cannot hybridize with common Brassica crops (e.g. *B. juncea, B. napus, B. rapa, or B. nigra*) preventing pollen-mediated gene flow into these high-value food crops (Canadian Food Inspection Agency, 2018).

Enthusiasm for the introduction of any uncommon crop should be balanced with ‘duty of care’ (Byrne and Stone, 2011). The ‘duty of care’ approach uses predictive risk assessments and risk management strategies to allow crop introduction while minimizing potential environmental risks such as new weeds, biological invasions, or deleterious gene flow events. Some biofuels crops have attracted special attention for their potential ecological risks (Andow and Zwahlen, 2006; Barney and DiTomaso, 2008; Byrne and Stone, 2011; Cousens, 2008; Craig and Degrassi, 2008; Raghu et al., 2010; Warwick et al., 2009). Concern has also been expressed regarding the effect of large-scale biofuel crops on non-target insects and pollinators (Stanley and Stout, 2013).

Camelina is not native to North America, and the species has been reported as an annual weed for over 100 years (Canadian Food Inspection Agency, 2017; Davis et al., 2011; Martin et al., 2017; USDA Plants Database: https://plants.usda.gov/core/profile?symbol=CASA2). Risk assessments for novel crop species and GE traits have focused on their potential direct and indirect impacts as weeds or invasive species. Negative impacts could involve damage to native species or natural communities, reduced ecosystem services, harmful changes to the physical environment (e.g. soil, water), or harm to agricultural production (Cousens, 2008; Keese et al., 2013; Warwick et al., 2009). In general, the GE traits of greatest concern are those that increase plant fitness (e.g. nitrogen use efficiency, salt tolerance), create persistent volunteers in agricultural fields (e.g. herbicide resistance), or promote invasion in natural areas (Beckie et al., 2010; Cousens, 2008; Keese et al., 2013; Warwick et al., 2009). A model for this type of risk analysis has been canola (*Brassica napus*), an annual oilseed crop that grows as a weed outside cultivation and is related to camelina (Beckie et al., 2003; Beckie et al., 2010; Schafer et al., 2011). Weed risk assessment protocols (WRAs) and trait-based models have been advanced as methods to identify potential risk (Cousens, 2008; Davis et al., 2011; Keese et al., 2013). However, WRAs have been limited to regions outside of the Northeastern US.

Plant gene flow is defined as a change in gene frequency due to the movement of pollen, seed, individuals, or groups of individuals, and the ‘duty of care’ concept can be extended to concerns about gene flow from GE crops to co-localized crops, weeds, or wild relatives (Andow and Zwahlen, 2006; Craig and Degrassi, 2008; Ellstrand et al., 1999; Warwick et al., 2009). Pollen-mediated gene flow within camelina fields occurs at relatively low rates and short distances, but the mechanism for pollen dispersal and distance for pollen dispersal are unknown (Julié-Galau et al., 2014; Seguin-Swartz et al., 2013; Walsh et al., 2015). While pollinating insects have been reported on camelina flowers (Eberle et al., 2015, Groeneveld et al., 2014), their effect on gene flow and seed set (yield) has not been defined.

The major research goals in this project were to: 1) determine if camelina could produce reasonable yields in the Northeastern US, 2) understand the competitive ability of camelina in agricultural weed communities for future weed risk assessments in the Northeastern US, 3) identify and quantify the pollinating insects visiting camelina flowers, and 4) test for pollen dispersal through wind and insects.

## 2. Materials and Methods

### 2.1 Camelina cultivation

*Camelina sativa* ‘SO-40’ (donated by Sustainable Oils, California, USA) was grown at the University of Connecticut Plant Science Research Farm in Storrs, Connecticut for three years (2014, 2015, 2016) in a 0.32-hectare plot located at 41°80’N, 72°23’W. The farm is within the Level III 59 Northeastern Coastal Zone and Level IV 59c Southern New England Coastal Plains and Hills ecoregion (US Environmental Protection Agency, https://archive.epa.gov/wed/ecoregions/web/html/new_eng_eco.html, Last accessed June 23, 2016). The plot had a 3-8% West to East slope with a Paxton and Montauk fine sandy loam soil type with a soil pH of 5.8 (http://websoilsurvey.sc.egov.usda.gov/App/WebSoilSurvey.aspx). A weather station at the research farm (http://newa.cornell.edu/, Storrs Research Farm) provided information about precipitation and temperature. Growing degree days (GDD) were calculated as previously described with a base temperature of 10° C (Gresch and Cermak, 2011). Urea fertilizer (46-0-0) was applied at 168 kg/hectare before seeding in 2014 and 2015; fertilizer (15-15-15) was applied at 643 kg/hectare in 2016 due to low phosphorus conditions. The amount of applied N was the same across all years. No herbicides, insecticides, or irrigation were used.

Camelina seed were sown on at 6.27 kg/hectare (May 6, 2014), 6.5 kg/hectare (May 5, 2015), or 7.7 kg/hectare (April 29, 2016). Laboratory tests showed about 95% seed germination each year. The experimental design included random placement of 20 subplots (0.5m x 0.5m) in the field using geospatial coordinates and a random number generator. Camelina development in subplots was observed 12-16 times between seeding and harvest with data recorded on: number of camelina plants/subplot, date of first open flower, date of last open flower, percent camelina plants with open flowers, and percent plants with siliques. The flowering stage was defined as a plant with at least one open flower; fruiting was defined as siliques present without any open flowers. Harvest occurred when 75% of plants had mature siliques and seed. The above-ground portions of camelina plants and weeds were removed from 10 subplots at harvest on July 22, 2014, and 20 subplots on July 23, 2015 and July 26, 2016. Camelina plants were dried and data collected on: plant biomass (gdw); number of intact, half, or missing siliques; number of seeds/plant; seed biomass/plant. Weeds were removed from each subplot, identified to species level, dried, and weighed.

### 2.2 Insect pollinators

Insect pollinators were studied in two experiments using either insect exclosures or sweepnet transects. Insect pollinators were defined as those insects with the ability to move pollen from plant to plant (Lee-Mäder et al., 2011). Five 1m x 1m x 1m cubes (hereafter called insect exclosures) were built from polyvinyl chloride (PVC) tubing and placed at equal distances across the field. The PVC cubes were covered with netting (Outback travel net, Mombasa Brand, Arlington, Texas). The insect exclosures were placed in the field just before crop flowering and removed just after flowering to minimize any slight effects of shading within the structure. At harvest, 20 camelina plants were collected from inside each insect exclosure and 20 plants were collected from adjacent control plots where insects had access to the camelina flowers. The control plots were approximately 1 m to the east of each exclosure to minimize the effect of the structures on insect activity. Camelina plants were analyzed for biomass, silique number, seed number/ silique, seed number/plant, and seed biomass.

Sweepnet transects were conducted to identify and quantify pollinating insects in 2014 and 2016. No sampling was conducted in 2015 due to crop failure. Four sweepnet transects were conducted twice per day (10:00 am and 3:00 pm) on seven days before, during, or after flowering (approximately 32-65 days after planting) when the weather was favorable for insect activity (no precipitation, little or no wind). Insects were collected using a sweepnet (BioQuip, catalog number 7312MS, Rancho Dominguez, California) passed across the top third of the camelina plants while walking in a “W” pattern (four diagonal transects) across the field. Insects were transferred to plastic bags, frozen, and sorted into functional groups. Insects were identified in the lab using a microscope and taxonomic keys (Borror et al., 1976; Gullan and Cranston, 2010; Lee-Mäder et al., 2011).

### 2.3 Pollen Sampling and Identification

Camelina pollen was collected the air using four rotorod boxes located at the center of the field and at three locations 9m beyond the edge of the field (west, north and east). The rotorod boxes were constructed by the author (C. Auer) and housed a DC motor attached to a rotating-arm impactor with two narrow plastic impaction surfaces oriented vertically (plastic rods) that spin in a circle (Muilenberg, 2003). The plastic rods (2.98 mm width x 2.54 cm length on leading side) sampled a volume of air ranging from 0.70 m^3^ – 0.76 m^3^ per hour depending on the angular velocity of each rotorod box (velocities varied from 2900 – 3170 rpm). The leading side of each rod was given a thin coating of silicone grease and replaced every 30 minutes. After field exposure, the rods were transported to the lab where the number of pollen was counted using an Olympus SZH10 microscope (Olympus, PA, USA) along a 1-cm section at the center of each rod. Pollen counts were converted into number of pollen/m^3^/hour with an adjustment for the speed of each individual rotorod box where the m^3^ refers to the volume of air sampled.

To determine in honey bees were transporting camelina pollen, 10 bees were randomly selected and frozen. The pair of legs bearing pollen baskets were placed in water and DNA extraction (kit name) yielded 15 ng/ml DNA. SSR markers developed for camelina (Manca et al, 2013) were used to amplify DNA sequences from samples including: pollen harvested from honey bee legs, pollen from anthers of SO-40 flowers (positive control), leaves of SO-40 plants, leaves of *Arabidopsis thaliana*, pollen collected from a hive in North Carolina, and a negative control (DI water). Results from fragment were analyzed using Principle coordinate analysis (PCoA).

### 2.4 Crop:Weed competition

Two experiments were used to measure the ability of camelina to compete against agricultural weeds in the absence of herbicides. In each of three years, weeds in the subplots were removed at the time of crop harvest and identified to species level, counted, dried, and weighed. A separate crop:weed experiment was initiated in 2016 when camelina plants reached the 4-5 true leaf stage. Random subplots (0.5 m × 0.5 m) were established with minimal hand-thinning to contain four different densities of *C. sativa* (10, 25, 50, or 100 individuals). Two control subplots were established: camelina plants only (approximately 100 individuals) or weeds only (all camelina removed). These six treatment groups were replicated three times for a total of 18 subplots. All subplots were managed to maintain the desired treatments and harvested on July 25, 2016 when the camelina siliques were mature. Data was collected on number of camelina plants, number of weeds in each species, camelina seed number, and dry biomass.

### 2.5 Statistical Methods

Statistics were conducted in Microsoft Office Excel (Microsoft, Inc., Mountain View, CA), SAS (version 9.1, SAS Institute, Inc., Cary, NC) and SigmaPlot (version 11.0, SYSTAT, Chicago, IL). The first step was to analyze the data for homogeneity of variances and normality using SAS. Analysis of data was conducted using paired t-tests and ANOVA (SAS PROC T-TEST, PROC ANOVA). Differences in crop development and weeds (cumulative growing degree days, number of days for development, and weed species) were analyzed using a Fisher’s LSD test (SAS PROC GLM). A paired t-test was used to examine abundance of insect taxa in 2014 and 2016, and seed biomass in the insect exclosure experiment (SAS PROC T-TEST). SigmaPlot was used for correlation analysis and Pearson’s correlation ‘r’. The weed competition experiment (2016) was analyzed using ANOVA and Fisher’s LSD. Margalef’s Diversity index and Simpson’s Dominance index were calculated for the weed communities. Margalef’s Diversity index was defined as *D_mg_* = (*S* – 1) **/** ln (*N*), where *S* is the number of species present and *N* is the total number of individuals in the community (Gamita, 2010, Margalef, 1958). Simpson’s Dominance index was defined as D = Σ {[*n_i_* (*n_i_* – 1)] **/** [*N* (*N* – 1)]}, where *n_i_* is the number of individuals in species *i,* and *N* is the total number of individuals in the community (Heip et al., 1998).

## 3. Results and Discussion

### 3.1 Crop Performance

Camelina ‘SO-40’ seed was planted in three years at 6.3-7.7 kg seed/ha and observations were recorded on number of days to germination, first true leaves, and flowering (Table 1). The number of days to germination and first true leaves was similar in 2014 and 2016, although the value for growing degree days (GDD) was much lower in 2016. In 2015, the number of days to seed germination and the end of flowering were delayed relative to 2014 and 2016 (Table 1). The values for GDD at harvest (80 or 88 days) were 1115, 1294, and 1367. This result was similar to the 1230 GDD reported for plots in Europe and Canada (Zanetti et al., 2017). Flowering in 2015 was asynchronous and spread over 31 days compared to just 20-23 days in other years (Figure 1). Only 18% of camelina plants flowered concurrently in 2015 compared to 90-94% in 2014 and 2016 (Figure 1).

**Figure 1.**
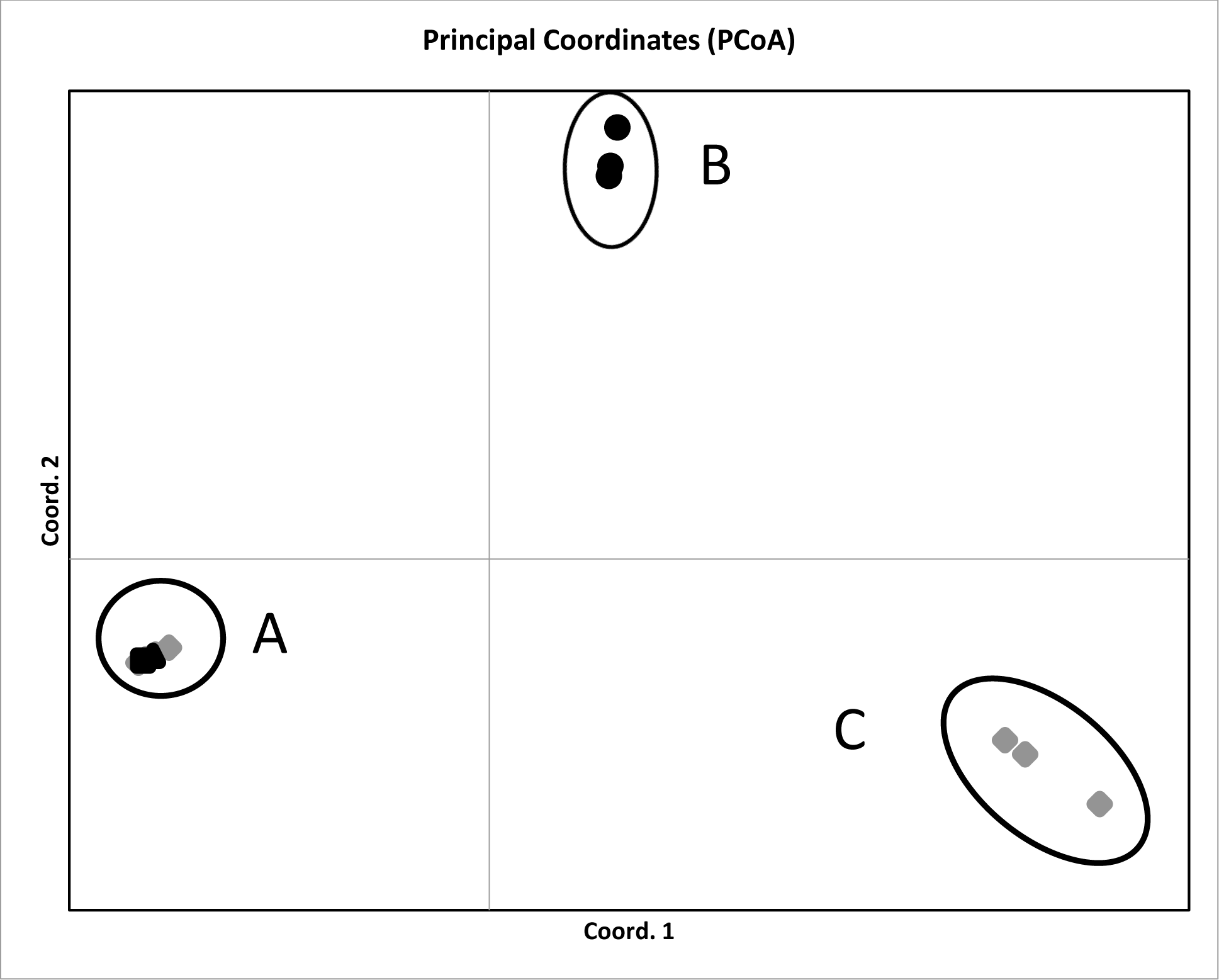
Principle coordinates analysis for SSR markers and fragment analysis for 5 samples. Circle A includes samples from *Camelina sativa* ‘SO-40’ leaves, *Camelina sativa* pollen, and pollen taken from honey bees in the camelina field. The outgroups shown are *Arabidopsis thaliana* leaves (circle B) and *Camelina alyssum* leaves (circle C).

**Table 1:** Observations of crop development at days after seeding and cumulative growing degree days (GDD). The timing of developmental events was based on observations in subplots (n = 20) except for the first observed seed germination which was based on observations across the entire field (n = 100). Mean and standard deviation are shown and letters indicate differences between years as determined by a paired t-test.

| **Developmental** | **2014** |  |  | **2015** |  |  | **2016** |  |  |
| --- | --- | --- | --- | --- | --- | --- | --- | --- | --- |
| **Event** | Mean #<br>of Days<br>after<br>Seeding | Range for<br>Days<br>after<br>Seeding | Mean<br>Cumulative<br>GDD | Mean #<br>of Days<br>after<br>Seeding | Range<br>for days<br>after<br>seeding | Mean<br>Cumulative<br>GDD | Mean #<br>of Days<br>after<br>Seeding | Minimum<br>and<br>Maximum<br>Days after<br>Seeding | Mean<br>Cumulative<br>GDD |
| **First observed**<br>**germination** | 4±2b | 2-7 | 11.3±3.4b | 19±13a | 2-33 | 54.4±24.4a | 6±3b | 2-10 | 0±0c |
| **First true**<br>**leaves** | 15±2a | 13-18 | 94.3±8.0b | 24±15a | 15-38 | 196.7±4.3a | 17±3a | 14-21 | 63.6±9.9c |
| **First anthesis** | 45±3a | 42-51 | 401.0±14.2b | 48±2a | 44-53 | 637.2±77.6a | 47±5a | 43-52 | 441.1±26.4b |
| **Peak anthesis**<br>**(>90%)** | 51±1a | 50-52 | 495.7±34.7b | ----- | ----- | ----- | 53±2a | 52-55 | 563.8±0a |
| **End of anthesis** | 60±3b | 57-65 | 889.8±103.6ab | 72±2a | 70-75 | 933.1±123.5a | 60±2b | 58-63 | 805.0±11.7b |
| **Harvest** | 80 | ----- | 1115.9 | 80 | ----- | 1292.4 | 88 | ----- | 1367.5 |

The number of camelina plants m^-2^ (stand density) changed during the growing season (Figure 2). Maximum stand densities were 403 plants m^-2^ (2014), 69 plants m^-2^ (2015), and 479 plants m^-2^ (2016). In 2014, the number of seedlings was highest at day 2 followed by decline of 50% at harvest. In 2016, the number of plants increased to days 8 - 17 and then declined by 41% to harvest. In these two years, self-thinning could have been due to competition between individual camelina plants, competition with annual weeds, or a combination of factors. A different pattern was observed in 2015 when camelina density reached a maximum at day 37 and then declined by 88% to harvest.

**Figure 2.**
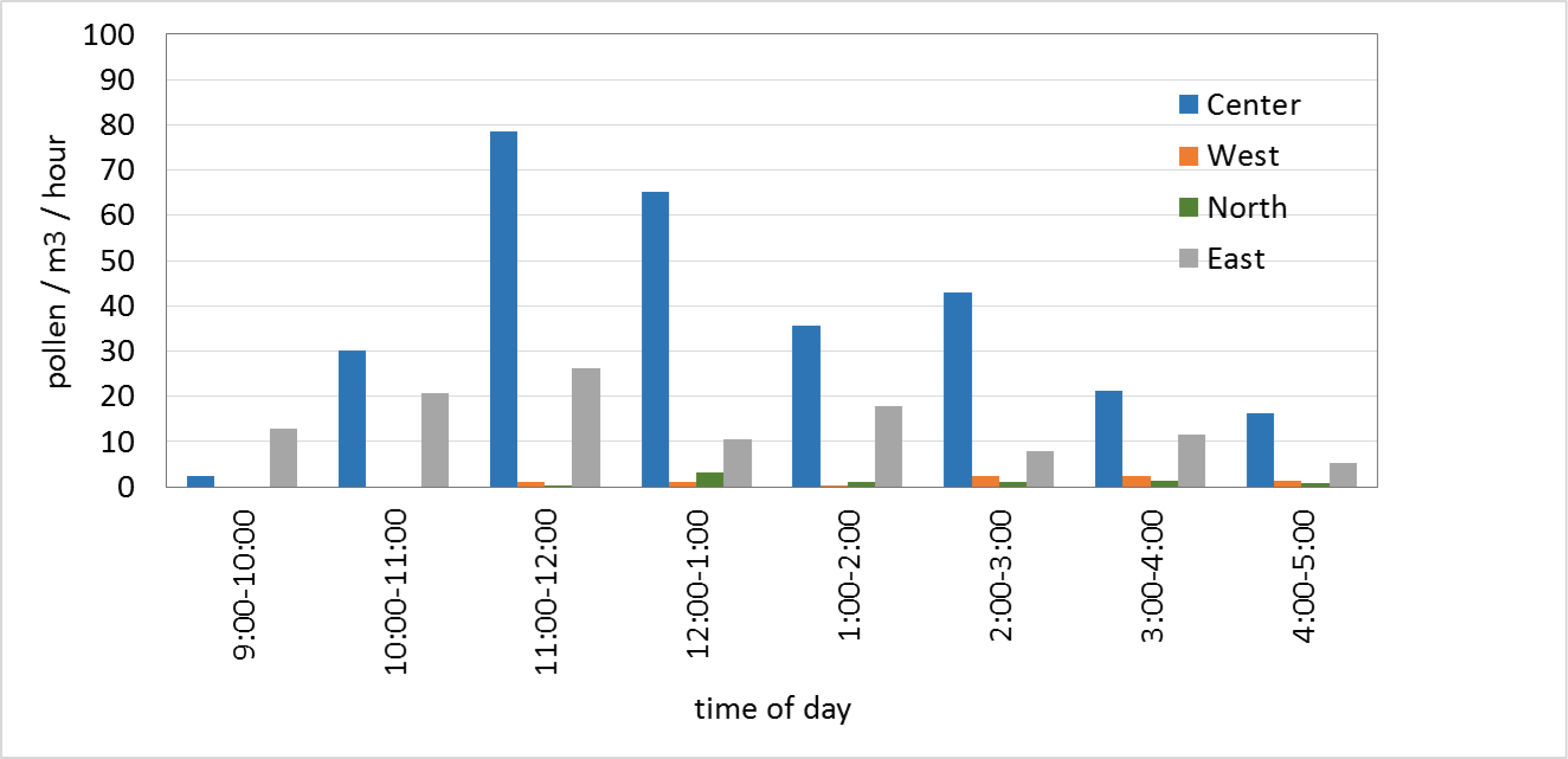
Camelina pollen concentrations over 8 hours when the field was at the peak of anthesis. Mean values are shown based on three days of sampling using rotorod boxes at the center of the camelina field and three locations from the edge of the field (9m to the west, north and east directions).

Camelina plant biomass, seed biomass, and number of siliques per plant were measured to assess plant performance and yield (Table 2). Again, very low values in 2015 demonstrated crop failure. Camelina plant biomass and seed biomass were higher in 2016 than in 2014. As reported in previous studies (Davis et al., 2013), seed biomass per plant was variable. This was due to plants that never flowered (∼ 6 - 10%), and plants that flowered but did not set seed. In addition, variations in soil conditions (e.g. moisture, fertility) may have contributed to the variation between the randomized subplots.

**Table 2:** Characteristics of harvested camelina plants. Mean values are shown with standard deviations. Letters indicate differences between years as determined by Fisher’s LSD. P<0.05.

|  | 2014 | 2015 | 2016 |
| --- | --- | --- | --- |
| **Plant biomass (mg)** | 640±560 b | 140±170 c | 1036±887 a |
| **Number Intact**<br>**Siliques/Plant** | 17.9±16.2 a | 3.6±5.7 c | 17.4±9.9 b |
| **Number Seeds/Plant** | 151.5±160.3 a | 7.9±17.8 b | ----- |
| **Seed Biomass/Plant (mg)** | 150±150 b | 10±20 c | 180±129 a |

Camelina seed biomass and plant density were used to calculate crop yield and 95% CI (Clewer and Scarisbrick, 2001). Yield was of 425.3 (286.4-564.2 CI) kg/ha in 2014 and 508.2 (379.4-637.0 CI) kg/hectare in 2016. These values fell within the range (372 - 1867 kg/hectare) reported for Montana (McVay and Lamb, 2004) and similar to values cited for ‘low yielding’ plots (500 kg/ha) in Colorado field trials (Jewett, 2013). The yield was lower than cultivars grown in South Dakota (1104-1158 kg/ha), Pennsylvania (1120 - 1681 kg/ha), Canada, or Europe (Hunter and Roth, 2010; Zanetti et al., 2017). While camelina yield in the Northeastern US was reasonable without herbicides, further studies are needed to optimize production variables (e.g. seed genetics, fertilizer rates, spring vs. fall planting).

Crop failure in 2015 prompted further analysis of environmental variables, precipitation, air temperature, and growing degree days (GDD). Figure 3 shows cumulative precipitation for each year compared to a 50-year average (1963-2013). Total rainfall in the three years was not significantly different at harvest (Figure 3). However, the first rainfall event in 2015 was 13 days after planting and the second rainfall did not occur until day 24. Rainfall at days 40 and 45 did not improve crop stand density which peaked at day 37 (Figure 2). Past research has emphasized camelina drought tolerance relative to canola and other oilseed crops (Bansal and Durrett, 2016; McVay and Lamb, 2008; Yang et al., 2016). However, other research has linked drought to poor stand establishment in South Dakota (Eberle et al., 2015), reduced yields in Western Canada (Gugel and Falk, 2006), and poor seed germination in Central Canada (Zanetti et al., 2017). Results in the Northeastern US support the conclusion that the timing of rainfall is critical to camelina seed germination, stand development, synchronous flowering, and good yield.

Temperature plays a role in seed germination and crop development, so cumulative GDD from planting (day 0) to harvest (80-88 days) was compared between years and with a 50-year mean (Figure 4). The value for GDD for each day in 2015 was significantly higher than 2014, 2016, or the 50-year mean (p < 0.001). Thus, it is reasonable to conclude that relatively high temperatures in 2015 contributed to poor crop development and very low yield.

This study in the Northeastern US showed that camelina in low-input conditions (fertilizer, no weed control, no irrigation) can produce approximately 500 kg/ha. Variations in weather between years showed that camelina seed requires rainfall early in the growing season for good germination and seedling development. Future research should optimize management practices to reduce self-thinning (∼41-50%) and increase yield. Camelina has high levels of cold hardiness (zones 1-9), and this hardiness trait would probably allow earlier planting dates in the Northeastern US when rainfall and soil moisture might be higher (Canadian Food Inspection Agency, 2017). Earlier planting dates might also allow better competition with weed populations.

### 3.2 Crop and Weed Interactions

Camelina was directly seeded into the field without herbicides or mechanical weed control. In this site, seven or eight common agricultural weed species were identified each year (Table 4). In addition, three weeds in the Brassica family were found at low abundance: shepherd’s-purse (*Capsella bursa-pastoris*), field pennycress (*Thlaspi arvense*), and pepper weed (*Lepidium latifolium*). Overall, the weed community was typical of agricultural weeds in the Northeastern US (Uva et al., 1997). The presence of shepherd’s purse was significant because of the potential for crop-to-weed gene flow. Previous studies have shown that camelina and shepard’s purse could be crossed to produce F1 seed, but the hybrid offspring were infertile (Julie-Galau et al., 2014; Martin et al., 2015). However, other ecotypes of shepard’s purse might be more interfertile.

Giant foxtail (*Setaria faberi*) was the most abundant weed in each year (Table 4). In the dry year of 2015, there were about 1000 foxtail plants m^-2^, although individual foxtail plants weighed less than in 2014 or 2016. Plots in 2014 had the lowest total number of weeds and total weed biomass. Plots in 2015 and 2016 had higher values for weed number and biomass, but these years were not significantly different from each other. This is surprising because camelina density was very low in 2015 compared to 2016 (Figure 2). In 2015, there was a slight negative correlation between the number of giant foxtail plants and the number of camelina plants (R = −0.626, n=20). Thus, it appears that the high temperature, low precipitation, and low camelina number in 2015 did not provide a major advantage to the weeds as measured by species diversity, number of individuals, or biomass. In 2014, there was a slight negative correlation between the number of ragweed plants m^-2^ and camelina plant biomass (Pearson’s correlation R= −0.633, n = 10). No other correlations were detected.

Margalef’s and Simpson’s Dominance indices were calculated to characterize diversity in the weed populations (Nkoa et al., 2015). Values for Margalef’s index was *d* = 0.983 (2014), *d* = 0.804 (2015) and *d* = 0.964 (2016). This showed low species richness and similarity between years. The values for Simpson’s index were D = 0.644 (2014), D = 0.700 (2015) and D = 0.371 (2016).

To further explore camelina’s ability to compete with agricultural weeds, randomized plots (n = 3) were established with treatments that modified the number of camelina plants (0-400 m^-2^) (Table 5). The two control treatments were 400 camelina without weeds (no weeds) and weeds only (no camelina). The plots of 400 camelina with or without weeds reflected the maximum observed crop stand density (403-479 plants m^-2^) in the field. Weed communities in each plot were allowed to develop naturally. Weed species were as reported (Table 4) except for the observation of three additional species at low abundance: shepherd’s-purse (*Capsella bursa-pastoris*), field pennycress (*Thlaspi arvense*), and pepper weed (*Lepidium latifolium*). The manipulation of camelina density had a significant effect on weed number (ANOVA p <.001), weed biomass (p <.001), and camelina seed yield (p <.001). Plots without camelina (weeds only) or with 40 camelina m^-2^ had similar numbers of weeds (∼1200 individuals m^-2^), but the weeds growing with 40 camelina m^-2^ had a reduction in biomass. Plots with 400 camelina m^-2^ had fewer weeds and lower weed biomass compared to plots with 100 camelina m^-2^. Camelina seed yield was equal in plots where 400 camelina grew without weeds or with weeds (∼585 weeds m^-2^) leading to the conclusion that camelina competed successfully with weeds in the conditions tested. Seed harvests from plots with 400 camelina m^-2^ gave an estimated yield of 1006 kg ha^-1^. Although this experiment was limited in scale, the results suggested that yields in the Northeast could approach the highest values reported for other regions (Zanetti et al., 2017).

This study showed that spring-seeded camelina can compete with agricultural weeds of the Northeastern US. Other studies have reported similar results. When used as a model weed, camelina reduced faba bean (*Vicia faba*) yield by 6-35% due to weed competition (Ghaouti et al., 2016). Another study showed that camelina suppressed annual weeds when planted between rows of field peas (Saucke and Ackermann, 2006). In contrast, a Weed Risk Assessment (WRA) study in Montana suggested that camelina was unlikely to become a serious weed (Davis et al., 2011). In that study, a qualitative WRA approach prohibited introduction of conventional (non-GE) camelina in the US, but field experiments and modeling from fall-seeded plots treated with three herbicides showed that conventional camelina was unlikely to become a serious weed in Montana (Davis et al., 2011). A greenhouse study using a species replacement design indicated that camelina was less competitive than spring canola against cheatgrass (*Bromus tectorum*) (Davis et al., 2013). Although not studied directly, annual weeds in the camelina fields may have contributed to pollinator abundance and diversity. There is increasing interest in managing agricultural weeds to provide resources (e.g. overwintering habitat, food) for pollinators. Further research is needed to determine the variables that affect camelina as a potential plant pest risk in agricultural fields or unmanaged areas.

### 3.3 Insect Pollinators and Camelina Pollen Dispersal

Insect pollinators on camelina flowers were captured and classified to their family or lower taxonomic rank when possible. The number of pollinators in the field changed over time with a temporal association between the maximum number of pollinators and the peak of camelina flowering (Figure 5). In 2014, the peak of flowering (95% at 50 days after planting) correlated with the peak number of pollinators (110 individuals) captured in sweepnet transects (Pearson correlation coefficient = 0.877). In 2016, 80% flowering at day 52 correlated with the largest number of pollinators (185 individuals) (Pearson correlation coefficient = 0.738). The number of pollinators at day 48 (90% flowering) was somewhat lower, but wind gusts up to 29 kph (18 mph) on that day may have reduced insect activity and abundance.

A diverse group of insect taxa was identified in four orders: Hymenoptera, Diptera, Coleoptera, and Lepidoptera (Table 3). The number of Hymenoptera (bees) collected in each year was higher than the other insect orders (Table 3). Very few Lepidoptera and Coleoptera were collected, but this may have been due to the sampling method and the behavior of these insects in the field (e.g. more difficult to capture than bees).

**Table 3.**
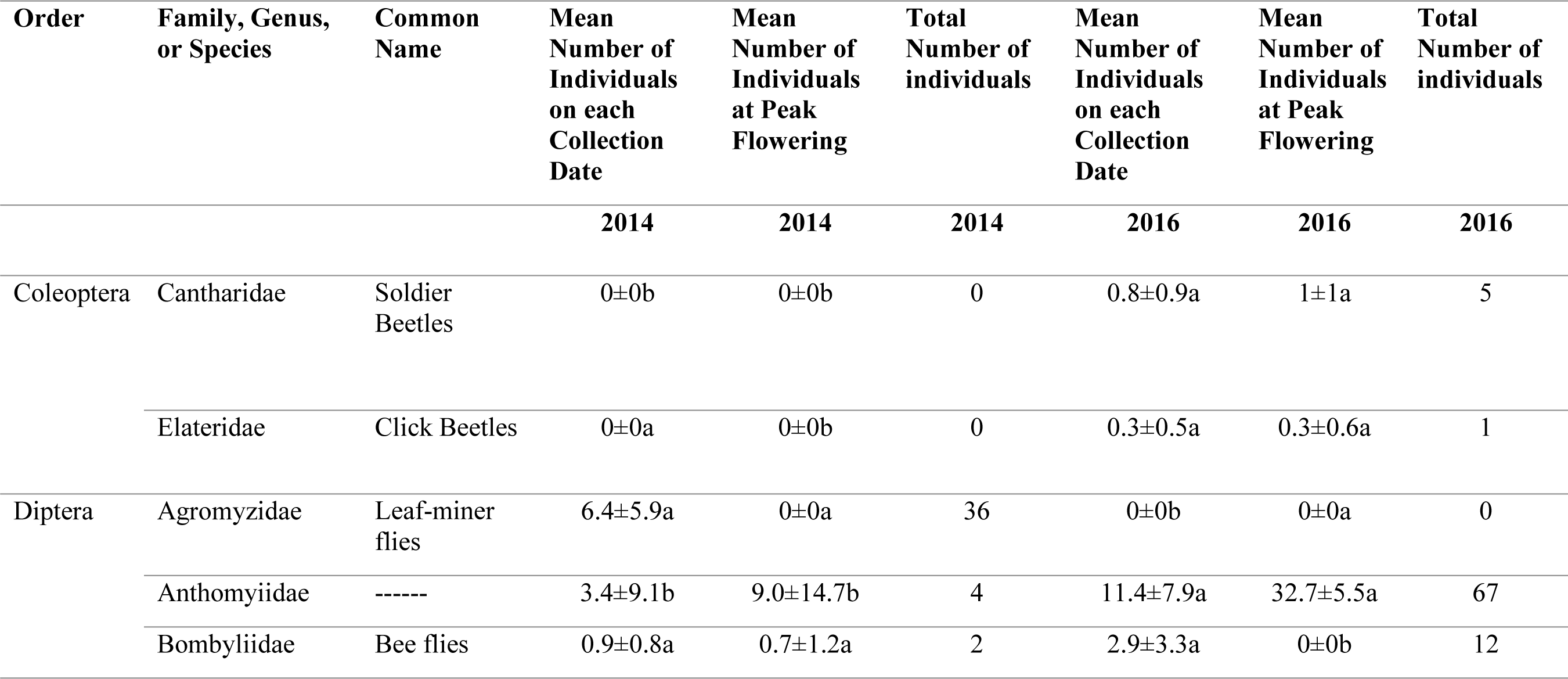
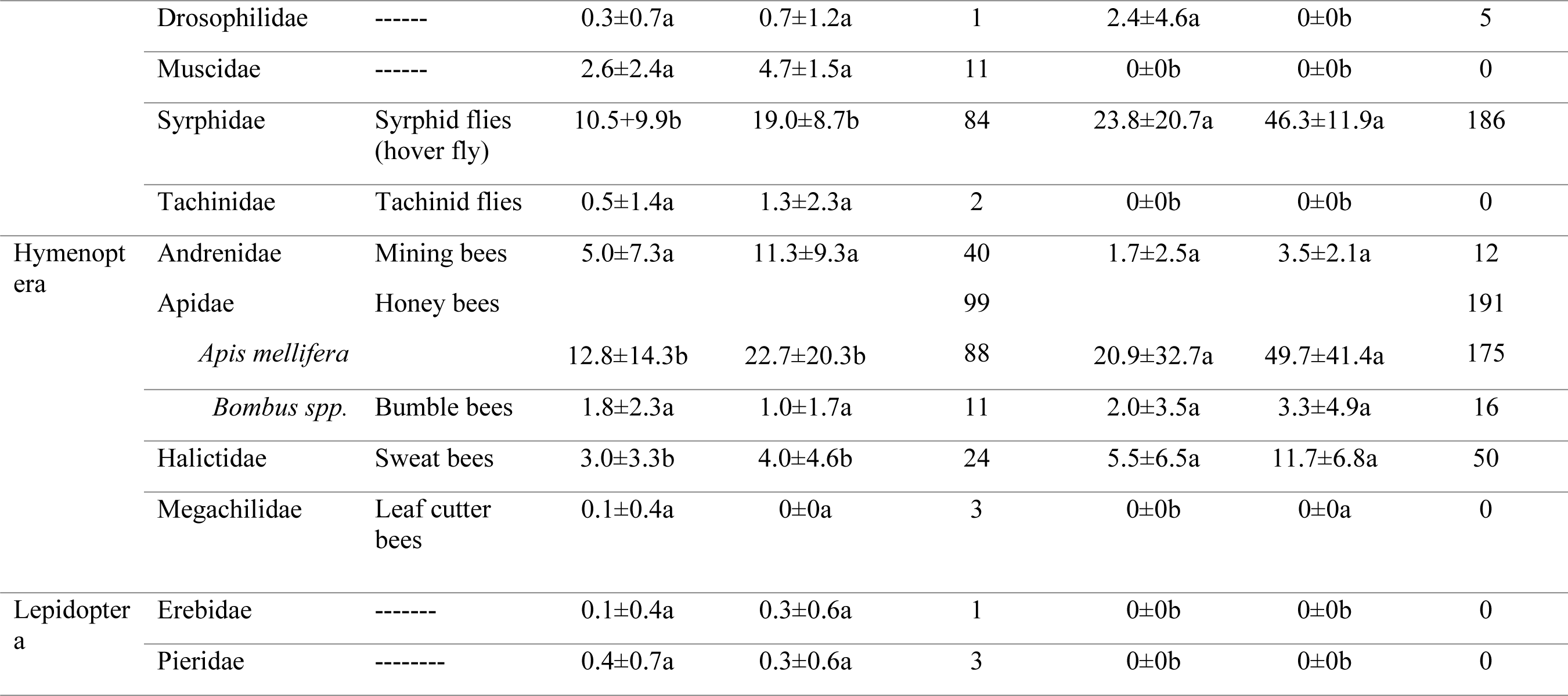
Insect taxa and abundance in camelina fields. The percentage shown for Apidae has been divided between honey bees and bumble bees (in parentheses). Mean and standard deviation are given for the number of individuals collected in transects on each collection date. The number of individuals at peak flowering is the number collected on the three days with the highest % camelina flowering (peak anthesis). Means with different letters show significant differences between years (2014 and 2016) based on a paired t-test.

**Table 4.**
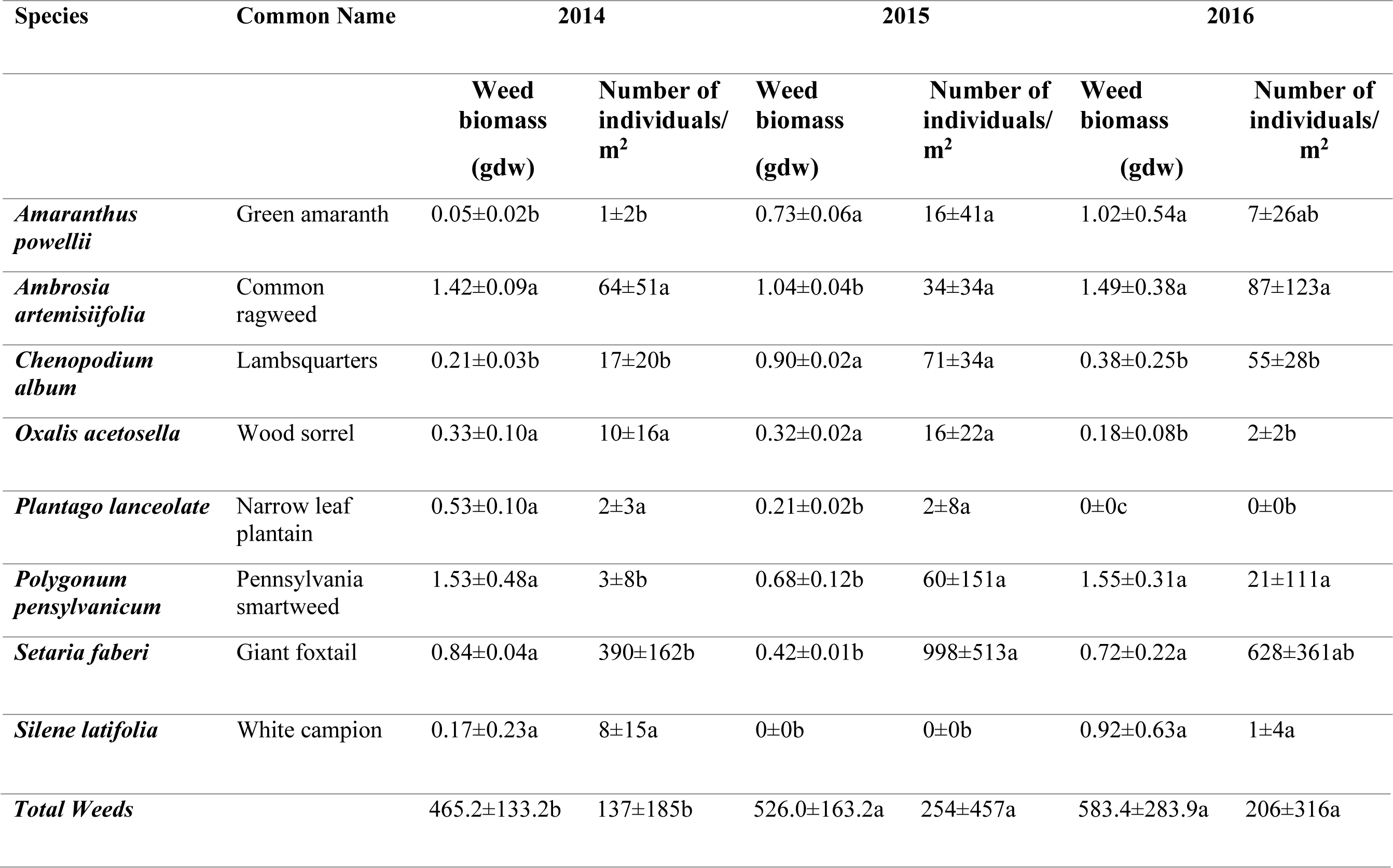
Agricultural weed community at crop harvest. Means and standard deviation are shown for subplots in each year. Letters indicate differences between years as determined by Fisher’s LSD. P<0.05.

**Table 5.**
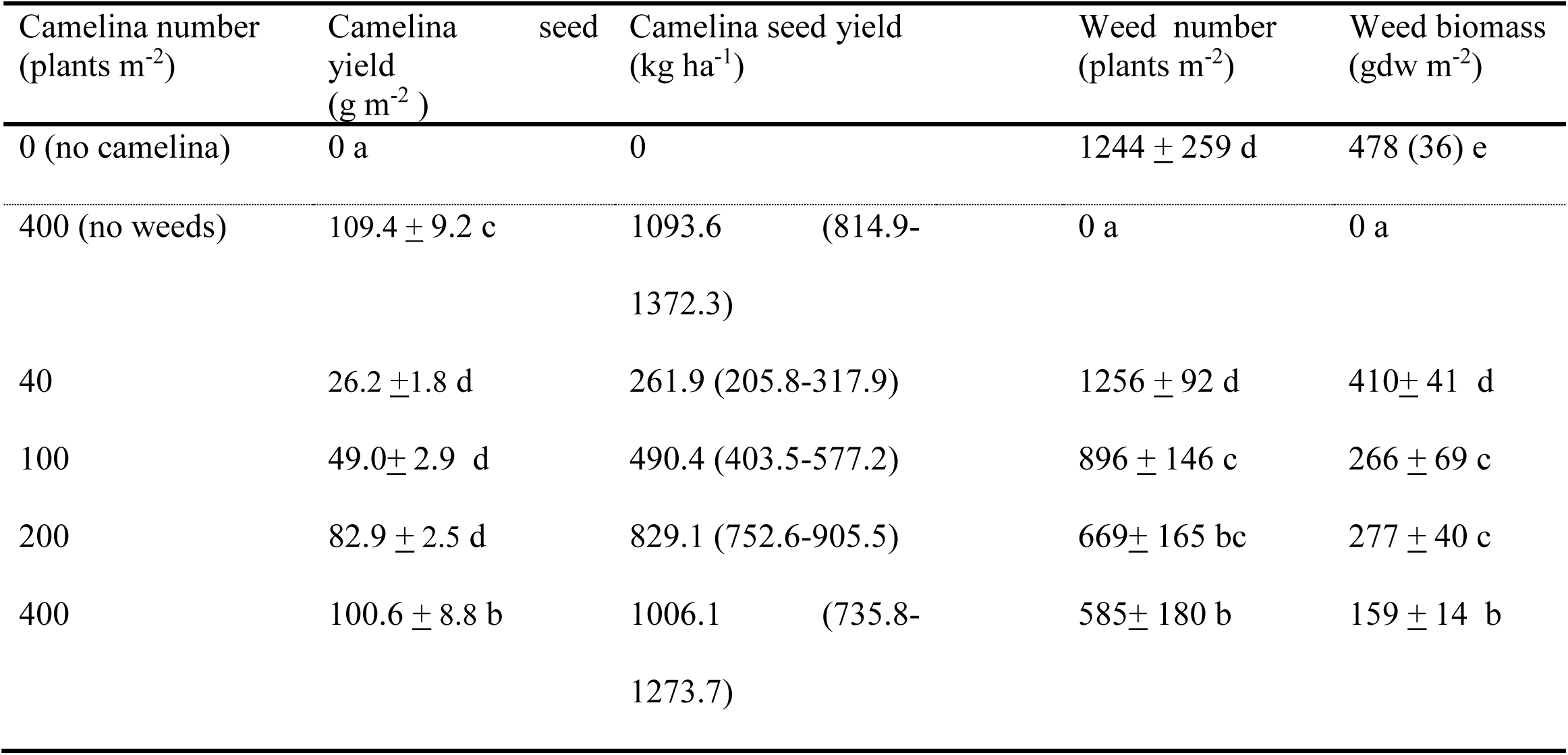
Effect of camelina number m^-2^ (stand density) on weed populations. Mean and standard variation are shown (n = 3) except for yield (kg ha^-1^) which shows mean and 95% confidence intervals in parenthesis. Letters within each column indicate differences according to Fisher’s protected LSD test at α=0.05.

The Hymenoptera included honey bees (*Apis melifera*), mining bees (Andrenidae), sweat bees (Halictidae), bumble bees (*Bombus* spp.) and leaf cutter bees (Megachilidae). Honey bees were the most abundant and leaf cutter bees were the least abundant (Table 3). Twice as many honey bees were collected in 2016 than 2014 (Table 3). The number of honey bees collected during the peak three days of crop flowering was about 2-fold higher in 2016 than 2014. However, honey bees were about the same percentage of all pollinators in both years (28% in 2014 and 33% in 2016).

An important question for gene flow is whether honey bees carry camelina pollen beyond the field borders. Pollen was taken from honey bee legs and other control samples for amplification with 15 SSR markers followed by fragment analysis and PCoA (Figure 1). Results showed that the five samples clustered in three groups: A) DNA samples from *Camelina sativa* ‘SO-40’ leaves, *Camelina sativa* pollen, and pollen taken from honey bees in the camelina field, B) DNA harvested from *Arabidopsis thaliana* leaves, and C) DNA from leaves of a congeneric species *Camelina alyssum*. These results confirm that honey bees not only visit camelina flowers, but also collect camelina pollen providing an avenue for pollen-mediated gene flow.

In the Diptera, syrphid flies (Syrphidae) were more abundant than other taxa, comprising 27% (2014) or 35% (2016) of all pollinators. Like the honey bees, the number of syrphid flies increased from 2014 to 2016 (Table 3). The larger number of pollinating insects in 2016 might have been due to the slightly higher density of camelina plants m^-2^ compared to 2014 (Figure 2). However, other factors cannot be excluded such as competition from other pollen and nectar resources in nearby weeds or native plants, pesticide applications, or weather conditions.

The timing, abundance, and diversity of pollinators indicated that camelina flowers are attractive and a good food resource in the agricultural landscape. This is an important factor in agricultural sustainability because many pollinators are in decline due to colony collapse disorder, agricultural intensification, habitat fragmentation and other complex factors (Eberle et al., 2015; Green et al., 2005; Potts et al., 2010; Stanley and Stout, 2013; Tilman et al., 1999). A previous study showed that camelina produced more nectar sugar than flowers of pennycress (*Thlaspi arvense*) or canola (*Brassica rapa*) (Eberle et al., 2015). In our study, honey bees made up a larger percent of total insect pollinators (28-33%) when compared to just 3% in a study in two Mid-Western states (Eberle et al., 2015). Conversely, camelina in Germany had a higher percent honey bees (47%) (Groeneveld and Klein, 2014).

Camelina also looks favorable when considering large-scale changes in land use for biofuels production. A study in Ireland showed that oilseed rape fields had more bumblebees than wheat or miscanthus grass (Stanley and Stout, 2013). This study also showed that all insect groups had higher species richness and abundance in the field margins where there were more flowers of non-crop plants (Stanley and Stout, 2013). Although not studied directly, annual weeds in the camelina fields may have contributed to pollinator abundance or diversity. There is increasing interest in managing agricultural weeds so that they provide resources (e.g. overwintering habitat, food) for bees and other pollinators (Food and Agriculture Organization, 2015). Further research is needed to determine if camelina could be managed in spring or fall plantings to optimize both yield and ecosystem services.

Insect exclosures were placed in the field to measure the effect of pollinators on seed yield. Camelina plants were exposed to pollinators or placed inside net cages to prevent insect visitation during flowering. An ANOVA showed no difference between years (p = 0.051), so yield data from 2014 and 2016 were combined (n=200 plants per treatment group). Seed biomass with pollinators was 0.29 ± 0.20 gdw/plant compared to 0.24 ± 0.16 gdw/plant without pollinators (inside cages). Thus, camelina seed yield increased modestly when insect pollinators were able to visit the flowers (Fisher’s LSD, p = 0.039 and Tukey’s multiple comparison). A similar study in Germany reported somewhat ambiguous results with a small yield increase for the cultivar ‘Celine’, but not for ‘Ligena’ or ‘Calena’ (Groeneveld and Klein, 2013).

While pollinators provide a crucial service to many crops and wild plants (Cresswell et al., 2002; Potts et al., 2010), these same insects could facilitate transgene movement to other crops, closely-related weeds, or native plants. A study on transgenic peanut plants reported that 50% of the pollen samples taken from bees included transgenic peanut pollen (Hu et al., 2015). A better understanding of bee visitation and pollen movement in camelina fields will be critical to designing effective biocontainment strategies for GE camelina, as well as promoting coexistence between farming systems.

### 3.3 Pollen dispersal by Wind

Camelina pollen was collected with 4 rotorod boxes over 8 hours on 3 days on days when the camelina fields were at the peak of anthesis. Pollen samples were captured in the center of the camelina field and 9 meters beyond the edge of the field in the west, east and north directions (Figure 1). The highest pollen concentration observed was 80 pollen/m^3^/hour in the center of the field in the middle of the day (11:00am - 1:00pm). This pollen cloud concentration was relatively low considering that >95% of the crop plants had open flowers, but consistent with the fact that camelina is primarily selfing (CFIA, 2017). Pollen concentrations collected beyond the field edge (9m) were considerably lower than in the center and ranged from 0 – 25 pollen/m^3^/hour. This if the first study demonstrating that small amounts of camelina pollen can be carried by the wind over relatively short distances.

## 4. Conclusions

Camelina is a relatively new oilseed crop with strong potential to contribute to the bioeconomy. However, introduction of this alternative crop must be balanced with ‘duty of care’ which asks for predictive ecological risk assessments and risk management strategies. This is especially important because camelina has traits commonly associated with weediness (e.g. rapid germination, short lifecycle), and it is already a minor weed in North America. This project showed that spring-sown camelina completed its lifecycle and produces 425-508 kg/ha with weed competition in the Northeastern US. This competitiveness could be helpful in organic production schemes where herbicides cannot be used. A variety of native and non-native Hymenoptera (bees) and Diptera (flies) were attracted to camelina flowers, and pollinator activity modestly increased seed yield. Experiments demonstrated for the first time that pollen dispersal could occur through honey bees or wind, although bee activity would likely be more significant for long-distance gene flow.

## Acknowledgements

This project was supported by Biotechnology Risk Assessment Grant award no. 2015-33522-24105 from the United States Department of Agriculture, National Institute of Food and Agriculture to CA. The authors wish to thank Dr. Ana Legrand (University of Connecticut) for her entomological expertise, and the farm crew for their assistance with crop planting and harvest.

